# A Machine Learning Approach to Predict Hypotensive Events in ICU Settings

**DOI:** 10.1101/794768

**Authors:** Mina Chookhachizadeh Moghadam, Ehsan Masoumi, Nader Bagherzadeh, Davinder Ramsingh, Guann-Pyng Li, Zeev N Kain

## Abstract

**Purpose:** Predicting hypotension well in advance provides physicians with enough time to respond with proper therapeutic measures. However, the real-time prediction of hypotension with high positive predictive value (PPV) is a challenge due to the dynamic changes in patients’ physiological status under the drug administration which is limiting the amount of useful data available for the algorithm.

**Methods:** To mimic real-time monitoring, we developed a machine learning algorithm that uses most of the available data points from patients’ record to train and test the algorithm. The algorithm predicts hypotension up to 30 minutes in advance based on only 5 minutes of patient’s physiological history. A novel evaluation method is proposed to assess the algorithm performance as a function of time at every timestamp within 30 minutes prior to hypotension. This evaluation approach provides statistical tools to find the best possible prediction window.

**Results:** During 181,000 minutes of monitoring of about 400 patients, the algorithm demonstrated 94% accuracy, 85% sensitivity and 96% specificity in predicting hypotension within 30 minutes of the events. A high PPV of 81% obtained and the algorithm predicted 80% of the events 25 minutes prior to their onsets. It was shown that choosing a classification threshold that maximizes the F1 score during the training phase contributes to a high PPV and sensitivity.

**Conclusion:** This study reveals the promising potential of the machine learning algorithms in real-time prediction of hypotensive events in ICU setting based on short-term physiological history.

## Introduction

The American College of Critical Care Medicine (ACCM) guidelines indicates that a Mean Arterial Pressure (MAP) of 60 to 65 mmHg is required for adequate organ perfusion [1]. Drop in the Blood Pressure (BP) below the limit for a prolonged period can lead to fatal consequences. This phenomenon is called arterial hypotension and frequently occurs in Intensive Care Units (ICUs) or Operating Rooms (ORs). New researches highlight the importance of predicting hypotensive events in hospital settings by providing evidences that correlate the occurrence of intraoperative or postoperative hypotensive events and various upcoming complications [2] [3] [4] [5] [6].

To promote the awareness among the scientific society, in 2009, Computers in Cardiology Competition challenged developers to predict hypotensive events using vital signals of 110 patients [7] [8] [9] [10] [11]. Following researches, including the study by Lee et.al [12] mainly adapted the methodology, rules and definitions laid out by this event [12] [13] [14] [15] [16] [17]. However, despite their significant contribution in showing the promising potential of machine learning in predicting hypotension, there are a few items that are missing or require further investigations. In particular:

- These studies defined hypotension as a drop in MAP below 65 mmHg for more than 30 minutes, which allows patients experience long period of low organ perfusion. No adequate clinical justification was provided for using this definition and its impact on ICU patients health status [12] [13] [14] [15] [16].
- These algorithms categorize the entire length of physiological record of each patient into hypotensive or non-hypotensive group. This approach of evaluating the algorithms only once per patient does not mimic the real-time monitoring applications during the entire ICU stay. For example, Lee et. al. [12] reported that by increasing the number of evaluation points per patient, the Positive Predictive Value (PPV) drops significantly from 66.5% to 13.6%, which leads to alarm fatigue among the hospital staff.
- The defined features required the data from a relatively long period of monitoring up to six hours [12] [13] [14] [15] [16], which impede the application of the algorithm in scenarios where only a short physiological history is available e.g. immediately after patients being admitted to the ICU.

To the best of authors’ knowledge, the research by Hatib et. al [17] is the only significant study in this field that addressed some of the above-mentioned challenges. However, the main target of the research was OR patients and the features were extracted from high-fidelity blood pressure waveforms using a propriety commercial software which neither is available to the public research community nor exists at all hospital settings. Also, the algorithm was evaluated at few specific predefined points of monitoring, which may artificially improve the reported accuracy by not considering the marginal data points. Further studies showed that the PPV of the algorithm can drastically drop down to 12.6% for the real-time monitoring scenarios in the OR settings [18].

Above mentioned shortcomings call for more in-depth analysis to judge the usefulness of the machine learning algorithms for prediction of hypotensive events in ICU. In particular, we believe there is a need to i) define the most clinically relevant definition of the hypotension for the ICU setting, ii) define the optimum number of required and readily available physiological signals to predict the events with a high sensitivity and PPV iii) the coverage of the algorithm across a large population of patient’s from multiple sites with a range of contextual information such as different medical history, hospitalization cause, administered drugs, etc., iv) train and test algorithms that considers changes in patients’ physiological state during monitoring rather than assuming each patient as one data point or only considering specific non-marginal data points, v) develop evaluation methods to better assess the algorithm performance and vi) to investigate the algorithm performance as a function of time to hypotension, which is required to find the best prediction window achievable by the adopted learning technique. Investigating all of the above items requires multiple focused studies each dedicated to specific item(s).

In this research, we attempted to address the last three challenges by developing a machine learning algorithm that mimics the real-time monitoring of the ICU patients. The proposed learning algorithm introduces a labeling approach that uses the majority of the data points from the patient’s physiological records to train and test the algorithm. It also requires only 5 minutes of prior physiological data to make the prediction at each data point within 30 minutes of the events’ onset. Because of the proposed labeling approach, we were able to evaluate the algorithm performance i) in a close to real-time monitoring scenario that includes all positive and negative points in the test sets and ii) as a function of time to event during 30 minutes prior to hypotension. These evaluation approaches help developers to determine the best prediction window that meets the algorithm performance objectives.

## Materials and Methods

### Problem Definition and Labelling Approach

The onset of a hypotensive event is tagged when the MAP drops below 65 mmHg in the next 30 minutes for at least 90% of the time i.e. 27 minutes in 30 minutes. We adopted this definition of hypotension solely to benchmark our results along with other available literature on using machine learning methods to predict hypotensive events in ICU units [12] [13] [14] [15] [16].

Figure 1 shows the data labeling scheme proposed in this study. Each data point of the physiological time series is categorized into positive (red), negative (green), or buffer (grey) zone. Positive points are defined as all the data points up to 30 minutes prior to the event onset. Negative points are defined as any point with MAP greater than 75 mmHg located at 40 minutes before or 20 minutes after any hypotensive event. This definition helps to separate the positive and negative zones [17]. Any data point that does not match these two criteria belongs to a buffer zone and is not used to train or test the algorithm. More than 70% of these data points in our dataset occur in the first 20 minutes after hypotension in which the patient’s physiological states are unstable due to the medical interventions such as drug administration. Although this is a common but considerable limitation that has been identified and investigated for the first time in this study, in practice the medical staff tend to monitor patients more closely in such periods of hospitalization. This helps to reduce the impact of the gray zone on the monitoring outcomes. Furthermore, as will be discussed later, other techniques can be used to reduce the size of buffer zones. Figure 2 shows the building blocks of the proposed algorithm, which will be discussed in detail in the following sections.

**Figure 1:**
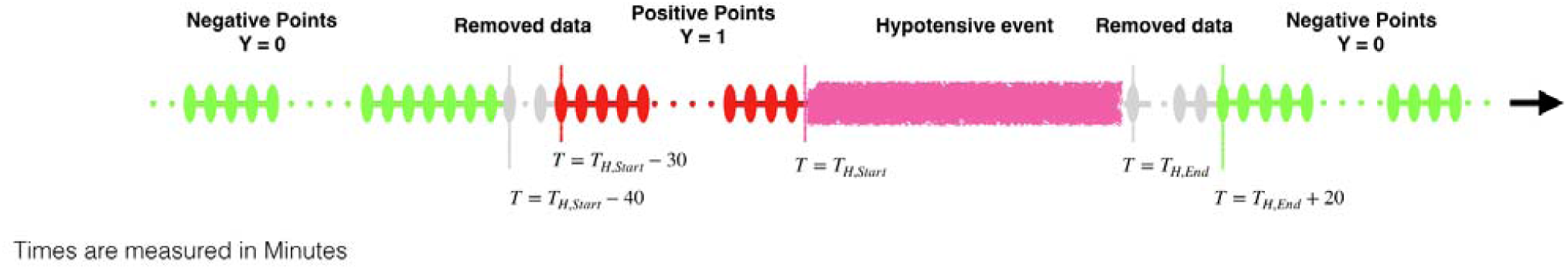
The data labeling approach. Data points up to 30 minutes before the onset of a hypotensive event labeled as positive (y=1). Any points 40 minutes before or 20 minutes after any hypotensive event labeled as negative (y=0). Gray points are not considered in the learning process.

**Figure 2:**
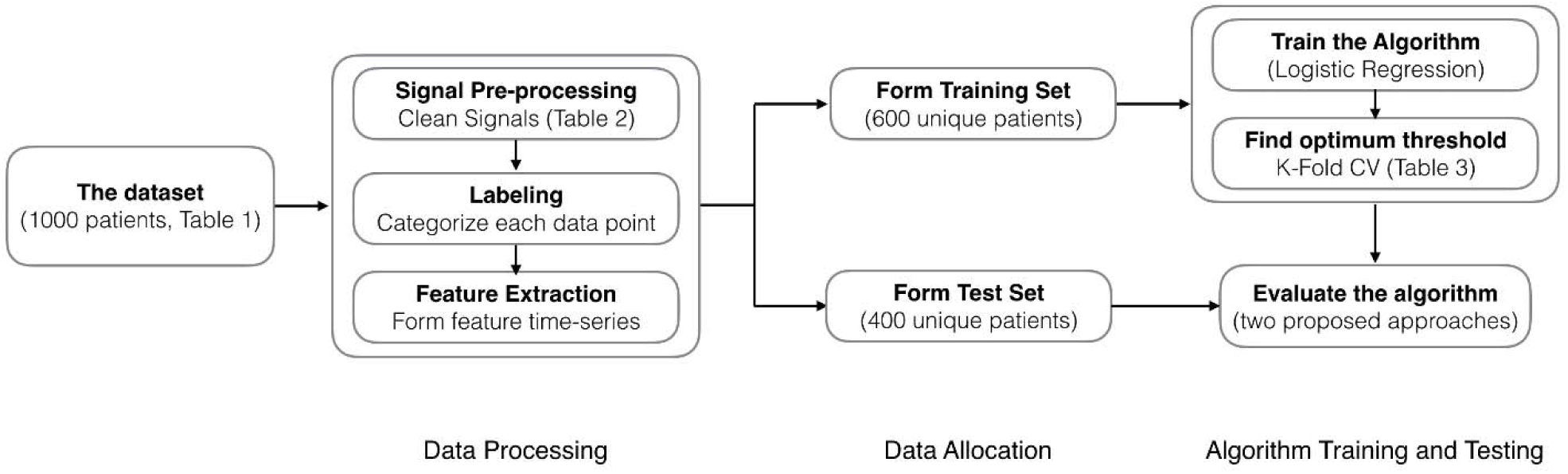
The overall flow of the proposed machine learning algorithm. The data from 1000 patients are going through the data processing steps from which 400 patients are randomly selected for the testing purpose and the remainings are used to train the algorithm. In the training phase, a 5-flod cross-validation is applied to find the optimum learning parameters which are used to evaluate the algorithm on the test set.

### Record Selection and Data Preprocessing

We randomly selected 1000 patient records from MIMIC III [19] [20] database version 1.0, released on August 2017, which have all the physiological signals of interest. Table 1 shows the patients’ demographics including age, gender, Body Surface Area (BSA) as well as the top reason of hospitalization for the selected patients. It can be seen that the majority of the patients have been admitted to the emergency room prior to ICU transfer. The numerical database of MIMIC III contains minute-by-minute physiological data of ICU patients which are driven from physiological monitors. We then preprocess each signal to remove i) spikes, where the variation of the signal value is more than 25% Of the baseline in one-minute period and ii) segments with values outside of the clinical range listed in

**Table 1:** Patients’ demographics including age, gender and reason of hospitalization.

| Number of Patients<br>(n) | Sex<br>(male) | Age<br>(year) | Weight<br>(kg) | Height<br>(cm) | BSA<br>( | Top Admission<br>Type | Top Diagnosis at<br>Admission |
| --- | --- | --- | --- | --- | --- | --- | --- |
| 1000 | 604 | $65 \pm 14$ | $83 \pm 23$ | $169 \pm 10$ | $2.0 \pm 0.3$ | Emergency (680) | Coronary Artery Disease<br>(190 patients) |

. Then, we assume that the missing parts of the physiological signals can be interpolated if the signal(s) value were missed for less than 5 minutes, otherwise that period is removed.

### Feature Extraction

We used information from three sensor lines which output six physiological signals including ABP (Arterial Blood Pressure), HR (Heart Rate), Systolic Blood Pressure (Sys), Diastolic Blood Pressure (Dia), Resp (Respiration rate), and peripheral capillary oxygen saturation (SpO2) for feature extraction. We then derived five additional numerical signals including Pulse Pressure (PP), MAP, Cardiac Output (CO), MAP to HR ratio (MAP2HR), and the average of RR intervals on ECG time series (RR), as following:

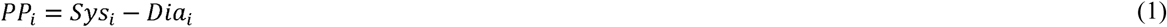

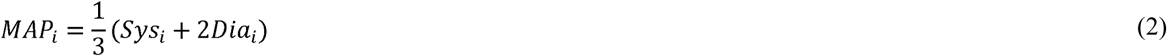

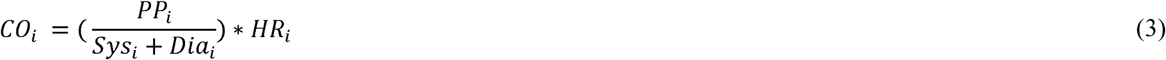

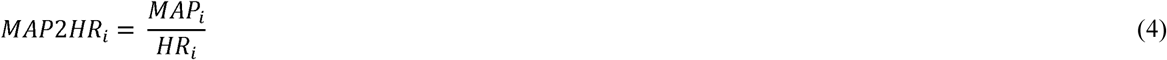

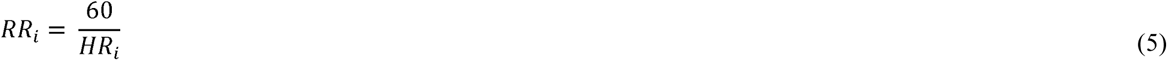

It should be noted that Equation (3) only provides a correlation with CO and does not quantify it [21]. These eleven time-series form our basic feature sets and represent the main physiological parameters at each one-minute interval. Furthermore, for each of these time series, we calculated short-term statistical features including, moving Mean and Standard Deviation (SD) for 5-minutes prior window as following:

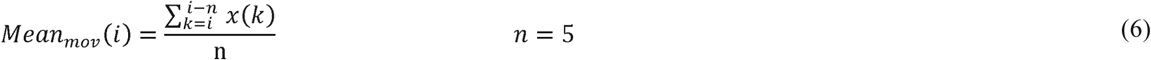

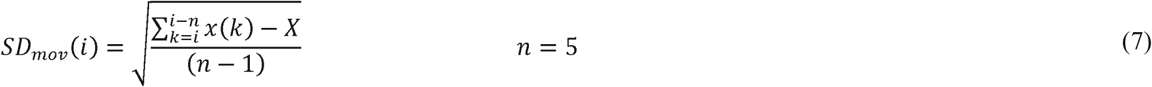

From the statistical point of view, using five data points to calculate the mean and standard deviation of a timeseries may not pass the normality test, however, it should be noted that equations (6) and (7) are just calculating features that their effectiveness in predicting hypotension should be analyzed at the training and testing phases.

### Data Allocation

We first identified the onset of hypotensive events which amounted to 2,710 total events in the entire dataset. The training set is then formed by allocating the entire records of 600 randomly picked patients, which their total number of hypotensive events amounts to around 65% of the total events i.e. 1,764 events. The remaining 400 records including about 946 hypotensive events formed the test set used for algorithm evaluation. Allocating the entirety of each record to either test or training set ensures that we do not overestimate the performance of the algorithm by using data points from patients in the training set to evaluate the algorithm. Each minute of monitoring either in the training or test set then is labeled as positive, negative or gray. To reduce the skewness, we down sampled the data points by randomly choosing 20% of negative data points during training of the algorithm as well as the testing phase. Table 3 summarizes the resulting data sets.

**Table 2:** Clinical range of physiological signals as well as central tendency and variation of the patients.

|  | BP<br>(mmHg) | Sys<br>(mmHg) | Dia<br>(mmHg) | HR<br>(beat/min) | SpO2<br>(%) | Resp<br>(breath/min) |
| --- | --- | --- | --- | --- | --- | --- |
| Min | 30 | 50 | 20 | 30 | 70 | 2 |
| Max | 150 | 220 | 103 | 180 | 100 | 35 |
| Central Tendency of the<br>studied Patients | 82 ± 17 | 123 ± 24 | 61 ± 13 | 85 ± 16 | 97 ± 2 | 18 ± 6 |

**Table 3:** Number of patients and data points in training and test sets.

|  | Number of records (patients) | Number of positive points | Number of negative points |
| --- | --- | --- | --- |
| Training and cross validation | 600 | 51,960 | 228,144 |
| Testing | 400 | 27,513 | 153,634 |
| All sets | 1000 | 79,473 | 381,778 |

### Data Labeling and Machine Learning Algorithm

We used the data labeling approach described previously to extract positive and negative data points for our classification algorithm. Throughout this study we developed in-house subroutines in MATLAB 2017b (MathWorks, Inc.) to train and test several classification algorithms including ridge logistic regression algorithm, variety of SVM algorithms and nearest neighbor algorithms with different kernels. In conjunction with these algorithms, we also investigated reducing the dimensionality of our feature space using Principal Component Analysis (PCA) retaining 95% of variance. We concluded that while most of these methods lead to similar outcomes, ridge logistic regression classifier yields better outcome. Logistic regression is a very popular method and its hypothesis function is defined as:

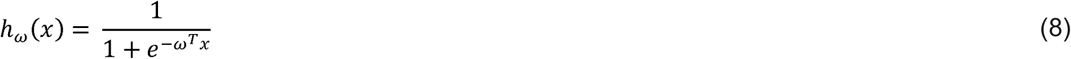

where ‘x’ is the independent variable (features) vector, ω the corresponding vector of coefficients, which are obtained by solving the following cost function:

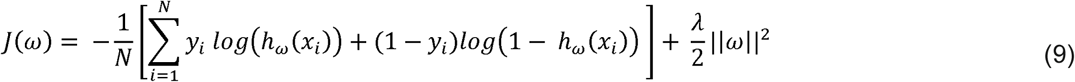

We used “fminunc” function of MATLAB to optimize the convex cost function of the ridge logistic regression algorithm with maximum of 400 iteration to converge.

### Optimizing the Classification Threshold

Logistic regression algorithm classifies a data point as positive if its output is bigger than a threshold. Bigger threshold values lead to a more conservative algorithm with a higher specificity and a lower sensitivity. Thus, the optimum threshold should be obtained for the best trade-off between these two antagonistic objectives. For example, Lee et. al defined the discriminatory threshold as the value that maximizes the summation of specificity and sensitivity [12] while Hatib et.al [17] found a threshold that minimizes the difference between these two metrics. However, these objective function does not necessarily yield to a higher PPV. Maximizing the F1 score is another popular approach among machine learning developers to optimize the algorithm performance to achieve a higher PPV and sensitivity on skewed datasets [22]. These three approaches are listed in Table 4 and to the best of authors knowledge there is no study to elaborate on the effect of different threshold optimization approaches on the prediction performance. In this study, we investigate how these three criteria affect the trained algorithm performance by first finding the thresholds which optimize each criterion on the 5-fold cross-validation set and then evaluating the algorithm performance on the test set using those thresholds.

**Table 4:**
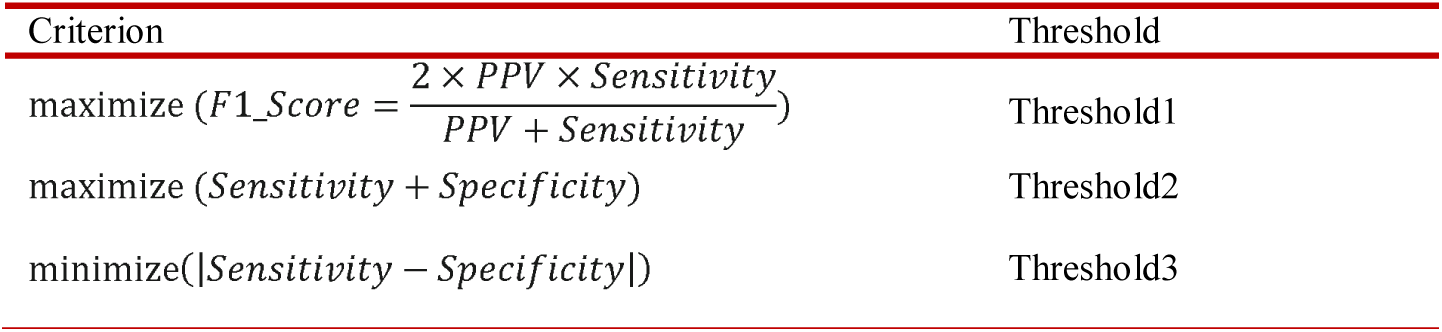
Different criteria to find the optimal threshold of the logistic regression algorithm.

### Evaluation Methods and Statistical Analysis

Training the algorithm using features extracted from minute-by-minute time series allows us to evaluate the algorithm performance in a real-time scenario. Various competing Figures of Merits (FOMs) including accuracy, sensitivity, specificity, PPV, NPV (negative predictive values), and F1 score of the test set were reported with 95% confidence. The overall performance of the algorithm is a trade-off between these FOMs.

### Continuous Evaluation Method: Real-time Monitoring

This method provides lumped performance of the algorithm for all data points available in the test set. A positive flag means there will be a hypotensive event within the next 30 minutes and a negative flag means there is no hypotensive event at least in the next 40 minutes. This evaluation method mimics the field application of ICU settings where the patient is unattended for some period of monitoring. To the best of the authors knowledge, this is the first time that a machine learning algorithm performance is evaluated in a scenario close to the real-time monitoring with a significant number of negative events, which in turn can severely affect the PPV.

### Discrete Evaluation Method: Selective Time Stamps to Hypotension

This novel approach extends the method used by Hatib et. al. [17] in which their algorithm was evaluated at only 15, 10, or 5 minutes prior to the hypotensive events and at the *midpoint* of 30 minutes intervals far from any hypotension. In this study, we calculated the predictive power of algorithm as a function of time to hypotension. To achieve this goal, we formed 30 sets of positive points located at the same time distances (one to 30 minutes) to the hypotensive events. Also, a set of representative negative points is created by picking *a random point* from each 30 minutes intervals located at least 40 minutes before and 20 minutes after any hypotensive event. Then, by evaluating the algorithm performance for the negative set as long as each of the 30 positive sets, we come up with the FOMs for the corresponding prediction window (timestamp). This approach provides a high resolution of the algorithm performance as we get closer to the events.

## Results

Figure 3 shows the Receiver Operating Characteristic curve (ROC) on the training set for a range of different thresholds used in logistic regression algorithm. The proposed model had an area under the curve of 0.93 (0.932-0.939 with 95% CI). The feature importance graph is also investigated (Figure 4) and shows that MAP2HR is the most prominent feature followed by RR.

**Figure 3:**
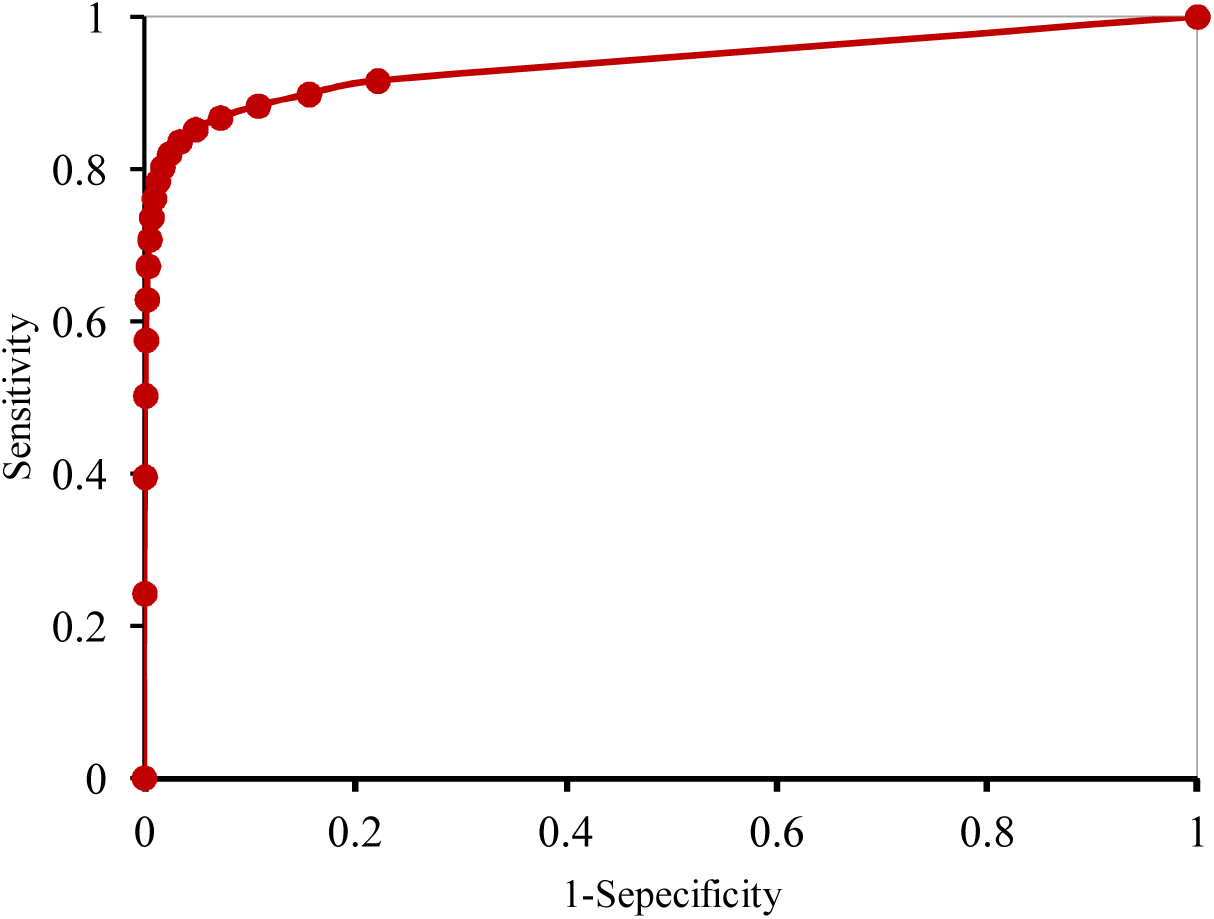
Receiver Operating Characteristic Curve (ROC) of the machine learning model for the prediction of hypotensive events in the training set. A high Area Under the Curve (AUC) of 0.93 (0.932-0.939 with 95% CI) represents a high potential of the algorithm to predict events correctly.

**Figure 4:**
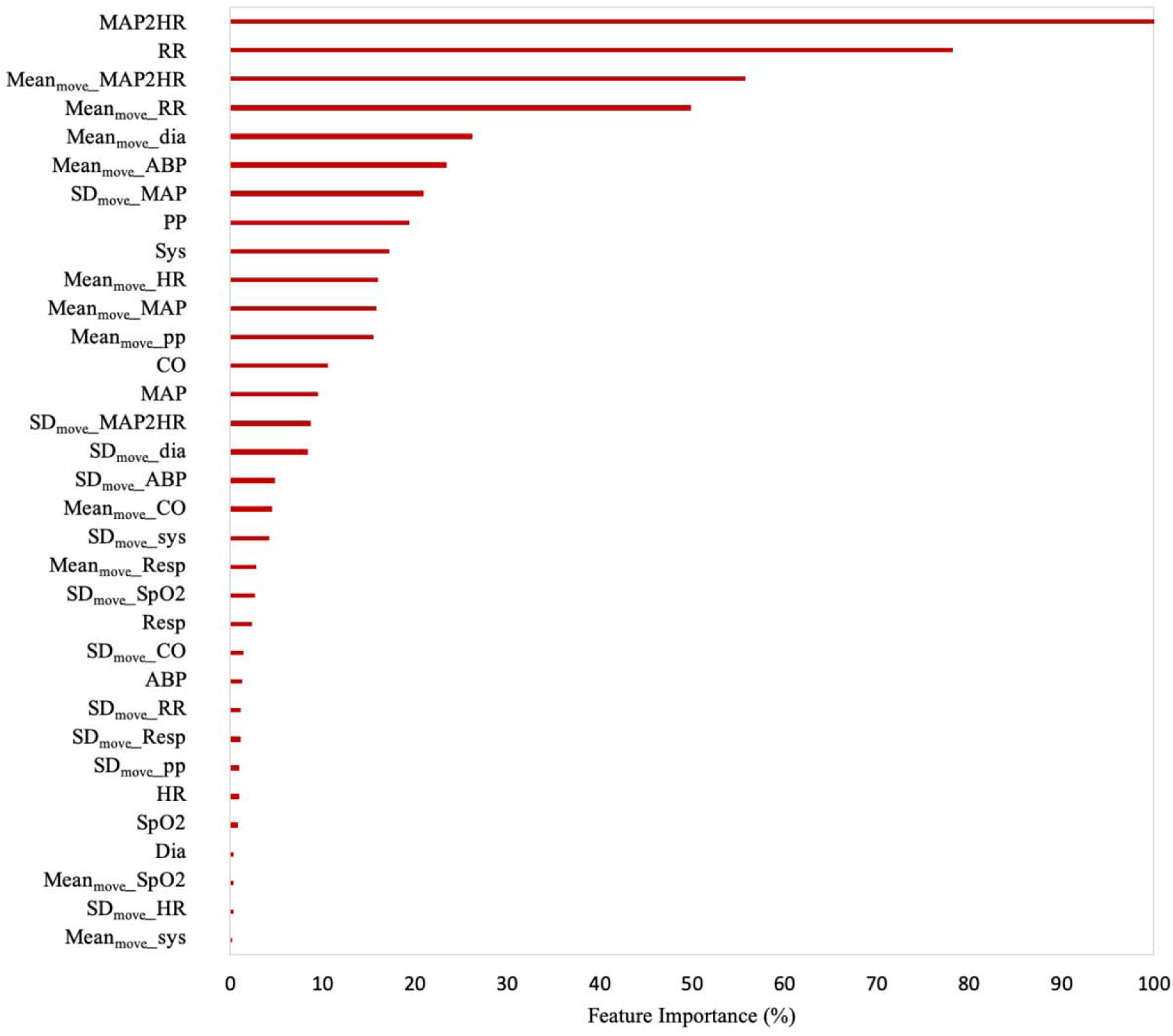
The importance of the features used to train the proposed machine learning algorithm. The greater the rate, the more essential the feature in the algorithm performance and its predictive power.

During the training phase, the threshold is optimized to apply the trained logistic regression algorithm on the test set. Table 5 shows the algorithm performance on the test set for three different threshold optimization schemes. During the real-time monitoring, the algorithm can predict hypotensive events within 30 minutes by 95% accuracy, 85% sensitivity and 96% specificity. Also, the model yields to an AUC of 0.92 (0.920-0.927 with 95% CI) on the test set.

**Table 5:** The algorithm overall predictive performance based on the real-time monitoring scenario, i.e. continuous evaluation method, for three optimization approach of Table 4. The results show that the first threshold optimization approach, which is based on optimizing the F1 score, leads to better results.

| Threshold optimization metric | Threshold | TP | TN | FP | FN | F1 (%) | Accuracy (%) | Sensitivity (%) | PPV (%) | Specificity (%) | NPV (%) |
| --- | --- | --- | --- | --- | --- | --- | --- | --- | --- | --- | --- |
| Threshold 1 | 0.45 | 23447 | 148234 | 5400 | 4066 | 83 | 95 | 85 | 81 | 96 | 97 |
| Threshold 2 | 0.3 | 24424 | 138508 | 15126 | 3089 | 73 | 90 | 89 | 62 | 90 | 97 |
| Threshold 3 | 0.2 | 24965 | 127795 | 25839 | 2548 | 64 | 84 | 91 | 50 | 83 | 98 |

Notably, Table 5 shows that the threshold optimization criterion significantly affects the algorithm predictive performance. Our investigation shows that the threshold that maximizes the F1 score on the cross-validation set leads to the best algorithm performance during the evaluation phase, while using the threshold that minimizes the difference between sensitivity and specificity leads to the lowest F1 score on the test set.

Figure 5 shows a graphical representation of the number of true positives and false negatives at each time stamp within 30 minutes prior to the hypotensive events. From the total of 7,130 tested negative points in discrete evaluation method, the algorithm only flagged 25 false positives. By analyzing the first flagging point of the events (Figure 5), on average the algorithm detects events 26 minutes prior to their onset, while 80% of them were predicted at least 25 minutes earlier. This information is valuable to determine the best predictive window and it is an outcome of the second proposed evaluation method.

**Figure 5:**
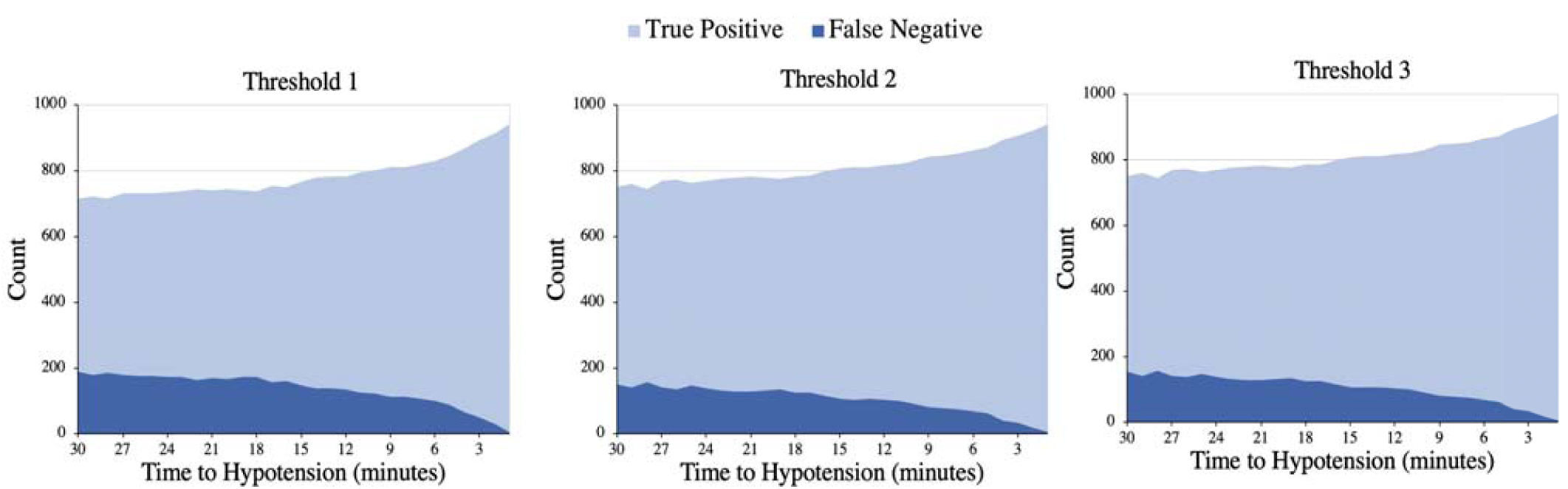
The number of true positives and false negatives at each time stamp within 30 minutes prior to the events for the three threshold optimization approaches.

Based on the results discussed above, we calculated the F1 score, sensitivity, and PPV of the algorithm as shown in Figure 6. Since the specificity is calculated based on negative data points, its value is not a function of time to hypotension. Hence, we did not include this parameter in Figure 6. As expected, Figure 6 indicates that the predictive power of the algorithm improves as we get closer to the hypotensive events and it is able to provide up to 96% PPV and 84% sensitivity, with 95% confidence, at 15 minutes prior the events while the specificity is evaluated as 99%. As a result of the evaluation approach presented in this work, Figure 6 shows that the sensitivity of the algorithm reaches to 80% mark, 24 minutes prior to the event for threshold1, however, high PPV ensures a reliable prediction throughout the entire 30 minutes prediction window.

**Figure 6:**
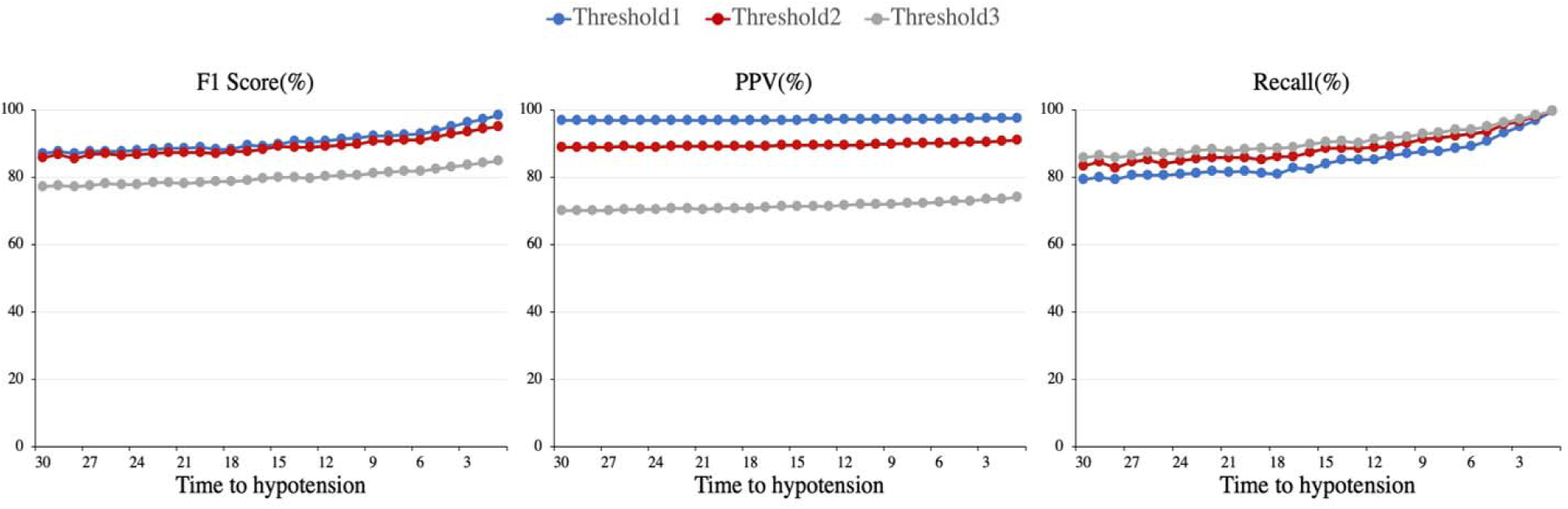
The predicative power of algorithm at each time stamp within 30 minutes before the hypotensive events for the three threshold optimization approaches.

## Discussion

Table 5 shows that throughout 181,000 minutes of monitoring of about 400 patients, our algorithm only flagged 5400 false positive which is reflected in 81% PPV. This significantly improves the experience of hospital staff compared to 13.6% PPV reported by Lee et. al. [12]. It should be noted that because of the high skewness of the dataset during the real-time monitoring, interpreting the algorithm performance based on the accuracy and PPV metrics is very difficult without providing the number of true positives and false negatives. However, since other researchers did not report this information, we only were able to compare our algorithm based on the common FOMs available in the literature.

It should be noted that in this study, we classified data points as positive or negative solemnly based on the output of the machine learning algorithm, which is far from a robust diagnostic platform with multiple layers of decision making. Developing a high-level risk assessment algorithm can further improve the prediction outcomes. For example, because of the high confidence of the algorithm in predicting negative points, in its simplest form such a risk assessment algorithm flags a positive event only if a number of specified consecutive data points are detected as positive. The detail implementation of this approach is out of the scope of this study.

Notably, both continuous and discrete evaluation methods show that maximizing F1 score leads to a significantly higher PPV, compared to the other two threshold optimization approaches. As previously described, this is an expected result since the F1 score optimization approach directly targets PPV while the other two approaches focus on finding a proper balance between sensitivity and specificity.

Interestingly for threshold2 and threshold3, evaluating the algorithm performance at distinct data points yields to significantly better results compared to the continuous evaluation method (Table 5 and Figure 6). One contributing factor to this discrepancy is that evaluating the algorithm at distinct positive or negative data points excludes the marginal ones where the algorithm is more prone to incorrect classification. This can also be a potential reason behind the lower PPV reported by [12] and [18] when considering marginal data points and highlights the importance of the proposed continuous evaluation method.

To benchmark our work along the other studies in the literature, we defined a hypotensive event as the drop of MAP below 65 mmHg for 90% of a 30 minutes window, which is not necessarily the best definition because of the relatively long period of low organ perfusion. We will address this limitation, in our next study by investigating the algorithm’s predictive performance for various definitions of hypotensive events.

Because of the short physiological history used to predict the hypotensive events, i.e. 5 minutes of prior data, this study hints the potential of using machine learning approaches to predict hypotensive events in the OR setting where the useful baseline data is limited due to the rapid changes in patient’s physiological state. However, since the algorithm has been trained and evaluated for the ICU patients, the scope of this study is limited only to the ICU units. Furthermore, we did not consider the reason for hospitalization, drug administration, type of surgical procedure, and other contextual data which may affect the algorithm performance in real OR setting. Addressing this challenge is the subject of our future works.

In this study to better train and evaluate the classifiers, we categorized some of the data points in gray zone, which is a deviation from field monitoring. To the best of author knowledge this is the first time that the coverage of the algorithms for predicting hypotension is discussed in the literature. While the labeling method used in this study has significantly reduced the gray data points compared to the other studies [12] [17], this limitation must be addressed before judging the application of machine learning algorithms in real time monitoring of hypotensive events. Developing multi-class classification algorithms, which classify each data point as positive, negative, or gray can be a potential solution for this limitation that is beyond the scope of this study. These algorithms can be combined with higher level risk management protocols to provide timely and precise prediction of the future events.

Moreover, in this study we used the continuous BP signal which is not available at all ICU bed sides because of its invasive nature. This limits the availability of the proposed method for all hospital settings. Training and evaluating a predictive algorithm based on only non-invasively measured physiological signals is a potential solution to address this issue. Also, because of its availability to the research community, we only used publicly available MIMIC III database. This is a major limitation of this study as expanding the trained algorithm to multiple hospitals with different protocols may affect the reported outcome.

## Conclusion

To study the potential of using machine learning algorithms to predict hypotensive events in ICU settings, in this research, we trained a logistic regression algorithm that uses only 5 minutes of the preceding physiological data to make prediction at each data point. ABP, ECG, Respiration Rate, and Oxygenation level time series were used to extract features. We proposed to label all available data points in these time series as positive, negative or gray zones. It was shown that compared to the other two optimization approaches used in literature to find the best classification threshold, maximizing the F1 score significantly improves the algorithm PPV. Overall, the algorithm was able to detect hypotensive events with 94% accuracy, 85% sensitivity, and 96% specificity. Furthermore, during 181,000 minutes of monitoring, the algorithm only flagged 5400 false positives (high PPV of 81%), which shows that it does not disturb the medical staff by false alarms. The algorithm sensitivity improved by getting closer to the event and passed the 85% mark at 22 minutes prior to the hypotensive event. The algorithm was able to detect 80% of the events at least 25 minutes earlier. Various limitation of the proposed method such as neglecting contextual data, single site study, gray zones, and making use of multiple sensor lines were discussed which are the subjects of our future studies.

